# Invasion triple trouble: Environmental fluctuations, fluctuation-adapted invaders and fluctuation-mal-adapted communities all govern invasion success

**DOI:** 10.1101/186254

**Authors:** Kati Saarinen, Leena Lindström, Tarmo Ketola

## Abstract

It has been suggested that climate change will lead to increased environmental fluctuations, which will undoubtedly have evolutionary consequences for all biota. For instance, fluctuations can directly increase the risk of invasions of alien species into new areas, as these species have repeatedly been proposed to benefit from disturbances. At the same time increased environmental fluctuations may also select for better invaders. However, selection by fluctuations may also influence the resistance of communities to invasions, which has rarely been tested. We tested eco-evolutionary dynamics of invasion with bacterial clones, evolved either in constant or fluctuating temperatures, and conducted experimental invasions in both conditions. We found clear evidence that ecological fluctuations, as well as adaptation to fluctuations by both the invader and community, all affected invasions, but played different roles at different stages of invasion. Ecological fluctuations clearly promoted invasions, especially into fluctuation mal-adapted communities. The evolutionary background of the invader clearly played a smaller role, and only at the later stages of invasion in the fluctuating environments. Our results indicate that climate change associated disturbances can directly increase the risk of invasions by altering ecological conditions during invasions, as well as via the evolution of both the invader and communities. Our experiment provides novel information on the complex consequences of climate change on invasions in general, and also charts risk factors associated with the spread of environmentally growing opportunistic pathogens.

**Data Accessibility:** Upon acceptance the data will be made publicly available.

**Statement of authorship:** Authors planned the questions and experiment together. KS conducted the laboratory work and wrote the first draft of the manuscript. TK conducted statistical analyses. All authors wrote the final version of the manuscript.

## Introduction

Current climate change scenarios predict that in addition to the increase in temperature, fluctuations in temperature and other environmental conditions are also increasing (Stocker *et al.* 2013) and creating selection pressures for biota. Some species benefit from these changes, and invasions and range expansions have been documented for many taxa (Parmesan & Yohe, 2003; Hickling *et al.* 2006; Bebber *et al.* 2013). In particular, fluctuations in environmental conditions might lead to the evolution of invasive genotypes (Lee & Gelembiuk, 2008) aiding species invasions. Thus, global climate change, with increased environmental fluctuations, could bring evolutionary and ecological problems to native fauna and flora: they must cope both with the direct changes caused by environmental fluctuations and with freshly-evolved invasive genotypes. This could be especially disastrous if native communities are mal-adapted to fluctuations.

One possible evolutionary explanation for the emergence of invasive species and genotypes is that they have evolved in a disturbed and fluctuating environment. It has been suggested that rapid fluctuations in particular, can select for traits that could promote invasion success, such as high population growth rate, plasticity and persistence (Levins, 1968; Turelli & Barton, 2004; Meyers *et al.* 2005; Lee & Gelembiuk, 2008). Climate change has been suggested to lead to increased extreme events (e.g. Stocker *et al.* 2013), but the pre-adaptive role of fluctuating environments on invasions has seldom been tested (Lee & Gelembiuk, 2008; Ketola *et al.* 2013). The literature on invasions has been centered on the evolutionary background of the invader, but the community’s properties, such as diversity and relatedness with the invader. can also influence invasion success (Davis, 2009). The evolutionary background of the invader, the community’s pre-adaptations to fluctuations, as well as prevailing conditions in general, could also dictate the success of an invader. If a community is adapted to fluctuations, environmental fluctuations should not cause repercussions in population size and hence benefit invaders.

Although it can be argued that all environments are potentially prone to invasions, significant variation exists in the sensitivity of different environments to invasions. Empirical evidence supports the theory that heterogeneous (both in space and time) and disturbed environments are more prone to invasions than stable environments (Burke & Grime, 1996; Davis *et al.* 2000; Davies *et al.* 2005; Melbourne *et al.* 2007; Liu *et al.* 2012). Disturbances might facilitate invasions by altering community composition, species competitive interactions, free resources, ecosystem processes, and propagule supply (Davis, 2009). Recent research has also recognized that invasions are interactions between the invader and the environment invaded, and as such the invasive traits might only make sense in certain environments (Burns, 2006; Facon *et al.* 2006; Mächler & Altermatt, 2012). Since disturbed environments can promote the evolution of invasive traits (Lee & Gelembiuk, 2008), studying the evolutionary background and pre-adaptations of the invader together with the effects of the current environment can also generate important information on the causes of invasions, especially in the context of current climate change (Facon *et al.* 2006). For example; will the increased fluctuations predispose communities to an increased risk of invasions, and will the fluctuation-adapted invaders invade such environments with even greater likelihood, as is suggested in the anthropogenically induced adaptation to invade – hypothesis (AIAI; Hufbauer *et al.* 2012)?

To test these key ideas about the ecological and evolutionary determinants of invasions (Facon *et al.* 2006), we used several species of microbes that had evolved for 2.5 months in two different kinds of environments; under constant temperature and under fluctuating temperature. These species and strains allowed us to make unique combinations of the invader (*Serratia marcescens*) and the community (*Pseudomonas chlororaphis, Enterobacter aerogenes* and *Leclercia adecarboxylata*) in which evolutionary adaptations either matched, or not, the environmental conditions during the invasions. With microbes it is straightforward to test general ecological and evolutionary theories, which are not amenable to testing with higher organisms (e.g. Buckling *et al.* 2009). Consequently, microbial experiments have also become more popular in invasion biology (Jiang & Morin, 2004; Jousset *et al.* 2011; Li & Stevens, 2012; Eisenhauer *et al.* 2013; Van Nevel *et al.* 2013; Amalfitano *et al.* 2015; Ma *et al.* 2015). However, the effects of environmental fluctuations, other than in resources, on invasion success (Davis *et al.* 2000; Li & Stevens, 2012) have rarely been tested (Kreyling *et al.* 2008). The invader species *S. marcescens* is an opportunistic pathogen capable of infecting various species such as plants, corals, nematodes, insects, fish and mammals (Grimont & Grimont, 1978; Flyg et al. 1980; Ketola et al. 2016 a). Hence, our experiment provides also an important test on the determinants of spread of pathogenic bacteria and diseases facing climate change induced fluctuations.

We hypothesized that if fluctuation in temperature is generally a driver of the evolution of invader genotypes (Lee & Gelembiuk, 2008; Ketola *et al.* 2013), experimental evolution in fluctuating environment should lead to bacterial clones that are more invasive. If instead fluctuations during invasion (i.e. on an ecological scale) lead to an overall improved invasion success it gives support to the idea that disturbance by various means enhances invasion success (see above for details, Burke & Grime, 1996; Davis *et al.* 2000; Davies *et al.* 2005; Melbourne *et al.* 2007; Liu *et al.* 2012). Moreover, if fluctuations increase the invasion success of fluctuation-adapted clones in particular, that would lend support to the anthropogenically induced adaptation to invade – hypothesis (AIAI; Hufbauer *et al.* 2012). This hypothesis postulates that human induced, often disturbed, habitats would allow for the invasion of those genotypes that have previously adapted to such conditions. Moreover, we expect that communities that have been evolving in fluctuating environments should be better in resisting invasions.

## Materials and methods

All the bacterial clones used in this experiment are from an evolution experiment (Saarinen, 2016), where we reared replicated bacterial populations (n = 10) of 9 different species separately (totalling 90 populations) at either a constant (30 °C) or thermally fluctuating regime (20 - 30 - 40 °C, at 2 h interval). The bacterial populations evolved in wells of a 100-well Bioscreen C^®^ (Growth curves Ltd, Helsinki, Finland) spectrophotometer plate in thermal cabinets ILP-12, Jeio Tech, Seoul, Korea). The experiment lasted 79 days, and every three days 40 *μ*l of culture was renewed to 0.4 ml of fresh Nutrient Broth medium (hereafter NB: 10 g of nutrient broth (Difco, Becton & Dickinson, Sparks, MD) and 1.25 g of yeast extract (Difco) in 1 l of sterile ddH20). After the experiment, we extracted 4 clones (i.e. colony forming units) from each bacterial population, utilizing dilution series plating, and stored the clones at −80 °C. This experiment yielded a total of 80 clones from each of the species (totalling 720 clones). *Serratia marcescens ssp. marcescens* (ATCC^®^ 13880™) was chosen as the invader because its ability to break DNA allowed easy recognition from other species using simple plating techniques (Ketola *et al.* 2016 b, Ketola *et al.* 2017). The 3 community species used in this study showed relatively high resistance against the invading *S. marcescens*, when reared together: *Pseudomonas chlororaphis* ATCC^®^ 17418™, *Enterobacter aerogenes* ATCC^®^ 13048™ and *Leclercia adecarboxylata* ATCC^®^ 23216™ (Ketola *et al.* 2017).

## Invasion experiment

As we were interested in the effects of invasion environment (constant vs. fluctuating temperature), evolutionary background of the invader (evolved in constant vs. fluctuating temperature) and evolutionary background of the community species (evolved in constant vs. fluctuating temperature) on the invasion success of *S. marcescens*, we set up an experiment testing all 3 effects together. We had 2 different thermal environments where the invasion took place: constant (30 °C) and rapidly fluctuating (2h 20 °C - 2h 30 °C - 2h 40 °C) and community members had either evolved in constant environment or in fluctuating environment. Moreover, in these very same conditions we tested how clones of *S. marcescens*, that had evolved in independent replicate populations (n = 10), either in fluctuating or constant temperatures, were able to successfully invade. The experiment was initiated by allowing the community species an assembly / establishment period prior to invasion (Fig. 1). Community species were mixtures of clones from 10 independently evolved populations evolved either in constant or in fluctuating environments (previously mixed and frozen at −80°C (1:1 in 80% glycerol)). After thawing, 20 μl of each species clone mix (totalling 60 μl) was pipetted into experimental 15 ml centrifuge tubes (Sarstedt, Numbrecht, Germany) filled with 6 ml of NB, and tubes were placed in thermal treatments: at constant 30 °C or at fluctuating 2h 20 °C - 2h 30 °C - 2h 40 °C (thermal cabinets: ILP-12, Jeio Tech, Seoul, Korea). Centrifuge tube caps were left loose to allow airflow. Throughout the experiment the cultures were stationary.

**Figure 1.**
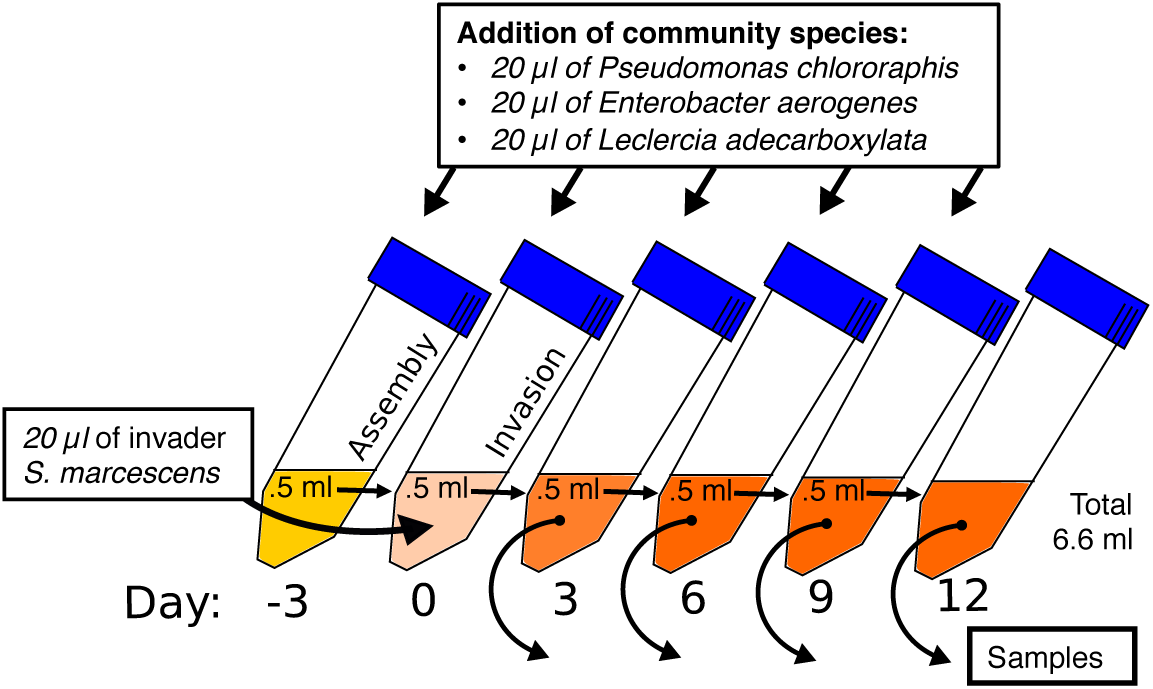
Overview of conduction of the invasion experiment, starting from the three day “assembly” period for the community, invasion three days later and renewal (to new tubes) and sampling (at −80°C in glycerol, for plating) every three days. This procedure was repeated with invaders adapted to fluctuating or constant temperature making invasion against community adapted to fluctuating or constant temperature in constant or fluctuating environment. Each of the eight combinations was replicated 10 times.

After the three day community assembly period we renewed the resources by pipetting a 500 *μ*l sample of each community into new tubes filled with 5.5 ml of NB. To maintain all community species in the community we supplemented cultures with 20 *μ*l of each community species clone mix (the clone mixes were also grown beforehand for 3 days at 30 °C, 60 *μ*l of bacteria in 6 ml of NB in 15 ml centrifuge tubes). The added clone mixes had always the same evolutionary background as the community species (Fig. 1).

After renewal and species supply, invasions were initiated to create the 8 different environment - evolutionary background combinations. The 20 (10 from fluctuating and 10 from constant environment) *S. marcescens* invader clones had been propagated beforehand for 2 days at 30 °C (60 *μ*l of bacteria in 6 ml of NB in 15 ml centrifuge tubes). To each community, we added 20 *μ*l of *S. marcescens* invader clones, which corresponds to 4 % of the renewed community (500 *μ*l) and is equivalent with the amount of external gene flow from each community species (see above). After which the communities, now with an invader, were placed in thermal chambers.

The communities were propagated for 12 days after the invasion, and we renewed the communities (as above), added the gene flow from the community species (as above) and froze samples of each community to −80 °C (final concentration of 40 % glycerol) every 3 days (3, 6, 9 and 12 days after the invasion). After the experiment we plated all the community samples from all the time points (3, 6, 9 and 12 days after invasion, altogether 320 samples). We used a standard dilution series technique to achieve a 10^5^-fold dilution that allowed the counting of separate colonies on agar plates. We plated the samples on DNase test agar with methyl green (Becton and Dickinson and Company, Sparks, MD; premade at Tammer-tutkan maljat, Tampere, Finland) that enabled the separation of *S. marcescens* from the community species (see: Ketola *et al.* 2016 b, Ketola *et al.* 2017). The fate of specific community species was not monitored during the invasion experiment, due to frequent supplementation of species.

## Data-analysis

To test if the experimental environment, evolutionary background of the invader, and evolutionary background of the community species had an effect on the invasion success of the invader (proportion of invaders), we used a generalized mixed model with a binomial error distribution and a logit link implemented in lme4 package in R. This kind of analysis is highly preferred instead of the linear models on arcsine square root transformed percentage data (Warton & Hui, 2011).

As explanatory variables we fitted experimental environment, evolutionary background of the invader, evolutionary background of the community species (constant vs. fluctuating in all cases) and all 2-, and 3- way interactions of the 3 fixed factors. Since we measured the same invader clone – recipient community combinations in two different invasion environments, we fitted the identity of the combination as a random factor to control for the non-independency of the observations. Time points (3, 6, 9 and 12 days after invasion) were analysed in separate models to facilitate the interpretation of the results.

## Results

After three days of the invasion, fluctuating environments supported higher invasions regardless of the invader’s or community’s evolutionary background (p<0.001, in all comparisons, Table 1 & 2). If the community had evolved in a constant environment it was more vulnerable to invasions in fluctuating environments than communities evolved in fluctuating environments (p<0.001). However, this effect was not visible if the invasion occurred in a constant environment (p=0.897, Table 1, Fig. 2a). Random effect of clone combination: σ^2^=0.3372, s.d.= 0.5807, t=0.130, p=0.898.

**Figure 2.**
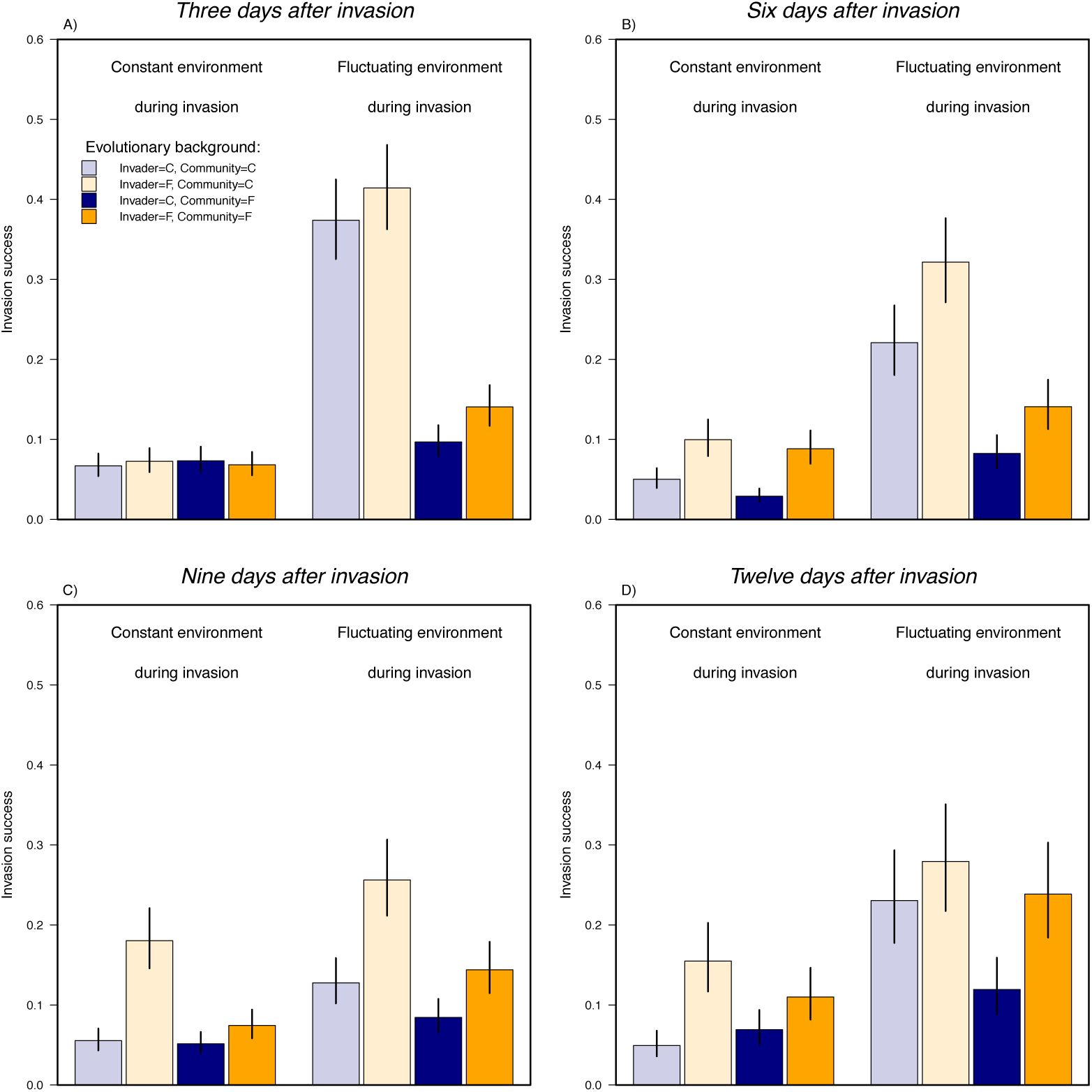
Proportion of *S. marcescens* colonies during invasion in bacterial communities **(A)** 3, **(B)** 6, **(C)** 9 and **(D)** 12 days after invasion, when invasion occurred either in constant or in fluctuating thermal conditions. Invaders that had evolved in the constant (blue) or in the fluctuating temperature (orange) invaded communities assembled from species evolved in the constant (lighter colours) or the fluctuating environments (darker colours). Figures correspond to the estimated marginal means and error bars reflect ± 1 standard error of the mean.

**Table 1.** Results of a mixed model exploring the effects of the evolutionary backgrounds of invader *S. marcescens* - bacteria and three species bacterial community, and environment during invasion, on invasion success 3, 6, 9 and 12 days after the experimental invasion.

|  | 3 days after invasion |  | 6 days after invasion |  | 9 days after invasion |  | 12 days after invasion |  |
| --- | --- | --- | --- | --- | --- | --- | --- | --- |
| | $\chi^2$ | p | $\chi^2$ | p | $\chi^2$ | p | $\chi^2$ | p |
| Invader Evolution | 0.4174 | 0.518 | 3.9587 | **0.0466** | 6.115 | **0.0134** | 2.2489 | 0.1337 |
| Community Evolution | 112.34 | **<0.001** | 95.647 | **<0.001** | 95.71 | **<0.001** | 20.143 | **<0.001** |
| Environment | 255.65 | **<0.001** | 282.91 | **<0.001** | 118.2 | **<0.001** | 246.24 | **<0.001** |
| Invader Evolution x Community Evolution | 0.1913 | 0.662 | 1.951 | 0.1625 | 19.97 | **<0.001** | 1.1421 | 0.2852 |
| Invader Evolution x Environment | 2.8184 | 0.093 | 4.9447 | **0.0262** | 2.943 | 0.0862 | 4.7472 | **0.0293** |
| Community Evolution x Environment | 87.055 | **<0.001** | 26.603 | **<0.001** | 0.006 | 0.94 | 12.527 | **0.0004** |
| Invader Evolution x Community Evolution x Environment | 1.5087 | 0.219 | 1.144 | 0.2848 | 7.091 | **0.0077** | 28.766 | **<0.001** |

**Table 2.** Post hoc comparisons of treatment level combinations. Only those factor interactions in which main test (Table 1) indicated significant differences are shown. Treatment combinations of environment, invader evolution and community evolution are indicated by the first three columns. Across column indicates over which treatment the test is done, followed by test statistics (Wald’s X^2^) and the significance of the comparison (p).

| Environment | Invader Evolution | Community Evolution | Across | $\chi^2$ | p |
| --- | --- | --- | --- | --- | --- |
| *3 days after the invasion* |  |  |  |  |  |
| Constant |  |  | Community Evol. | 320.277 | **<0.001** |
| Fluctuating |  |  | Community Evol. | 20.575 | **<0.001** |
|  |  | Constant | Environment | 0.017 | 0.897 |
|  |  | Fluctuating | Environment | 200.585 | **<0.001** |
| *6 days after the invasion* |  |  |  |  |  |
| Constant |  |  | Community Evol. | 277.481 | **<0.001** |
| Fluctuating |  |  | Community Evol. | 37.817 | **<0.001** |
|  |  | Constant | Environment | 7.783 | **0.005** |
|  |  | Fluctuating | Environment | 111.391 | **<0.001** |
| Constant |  |  | Invader Evolution | 112.094 | **<0.001** |
| Fluctuating |  |  | Invader Evolution | 106.467 | **<0.001** |
|  | Constant |  | Environment | 7.338 | **0.014** |
|  | Fluctuating |  | Environment | 2.627 | 0.105 |
| *9 days after the invasion* |  |  |  |  |  |
| Constant |  | Constant | Invader Evolution | 13.301 | **<0.001** |
| Constant |  | Fluctuating | Invader Evolution | 1.105 | 0.293 |
| Fluctuating |  | Constant | Invader Evolution | 5.792 | **0.048** |
| Fluctuating |  | Fluctuating | Invader Evolution | 2.606 | 0.213 |
| Constant | Constant |  | Community Evol. | 0.280 | 0.597 |
| Constant | Fluctuating |  | Community Evol. | 69.044 | **<0.001** |
| Fluctuating | Constant |  | Community Evol. | 12.193 | **0.001** |
| Fluctuating | Fluctuating |  | Community Evol. | 41.856 | **<0.001** |
|  | Constant | Constant | Environment | 61.522 | **<0.001** |
|  | Fluctuating | Constant | Environment | 18.877 | **<0.001** |
|  | Constant | Fluctuating | Environment | 11.529 | **0.001** |
|  | Fluctuating | Fluctuating | Environment | 35.142 | **<0.001** |
| *12 days after the invasion* |  |  |  |  |  |
| Constant |  | Constant | Invader Evolution | 7.260 | **0.028** |
| Constant |  | Fluctuating | Invader Evolution | 1.187 | 0.552 |
| Fluctuating |  | Constant | Invader Evolution | 0.306 | 0.580 |
| Fluctuating |  | Fluctuating | Invader Evolution | 3.219 | 0.218 |
| Constant | Constant |  | Community Evol. | 7.798 | **0.010** |
| Constant | Fluctuating |  | Community Evol. | 13.045 | **0.001** |
| Fluctuating | Constant |  | Community Evol. | 38.270 | **<0.001** |
| Fluctuating | Fluctuating |  | Community Evol. | 2.818 | 0.093 |
|  | Constant | Constant | Environment | 171.553 | **<0.001** |
|  | Fluctuating | Constant | Environment | 34.206 | **<0.001** |
|  | Constant | Fluctuating | Environment | 23.034 | **<0.001** |
|  | Fluctuating | Fluctuating | Environment | 67.418 | **<0.001** |

After six days of invasion, communities evolved in a constant environment were more vulnerable to invasion than communities evolved in fluctuating environments. This effect was bit more pronounced if the invasion occurred in a fluctuating environment (p<0.001) rather than a constant environment (p=0.005). Moreover, irrespective of the community’s evolution, fluctuating environments promoted higher invasion (all comparisons p<0.001, Table 2). After six days of invasion, an invader’s evolutionary background in a fluctuating environment promoted invasion success (Table 1, Fig. 2b). However, this was visible in constant environments during invasion (p=0.014) but not if invasion occurred in fluctuating environment (p=0.105). Random effect of clone combination: σ^2^= 0.5339, s.d.= 0.7307, t=0.164, p=0.872.

After nine days the invasion was stronger if the environment was fluctuating rather than constant during the invasion (Table 1, Fig. 2c in all comparisons the effect of environment was clear p<0.001, Table 2). If the community had evolved in a constant environment, invasion was stronger than if it had evolved in a fluctuating environment. The only exception to this pattern was if the invader had evolved in a constant environment and if the invasion occurred in constant environment (p=0.596, all other pairwise comparisons p<0.001, Table 2). Invader evolution in fluctuating environments was found to increase invasion success only if community species had evolved in a constant environment, and this result held in both invasion environments (constant: p<0.001, fluctuating: p=0.048, all other comparisons p>0.2, Table 2). Random effect of clone combination: σ^2^= 0.582, s.d.= 0.7629, t=0.171, p=0.866.

After twelve days fluctuating environment facilitated invasions regardless of the evolutionary background of the invader or the community (p<0.001, in all comparisons, Table 2). Invasion was also stronger if the community had evolved in a constant environment. However, when both the invader and the community had evolved in a fluctuating environment there was no difference in invasion success due to environment (p=0.09, all other comparisons within treatment combinations p<0.01). Invader’s evolution in fluctuating environments facilitated invasions only if environment was constant and if the community had evolved in a constant environment (p=0.028, all other comparisons within treatment combinations were clearly non significant, p>0.2, Table 2). Random effect of clone combination: σ ^2^= 1.005, s.d.= 1.003, t=0.224, p=0.825.

## Discussion

As climate change increases disturbances (Stocker *et al.* 2013) it is possible that invasions, range expansions, and evolution of invasive genotypes will become more common (Clements & Ditommaso, 2011; Bebber *et al.* 2013; Davis, 2009; Lee & Gelembiuk, 2008; Hufbauer *et al.* 2012; Ketola *et al.* 2013). We tested if evolution in a fluctuating environment, and fluctuations during invasions, affect the invasion probability of an environmentally growing opportunistic pathogen (*S. marcescens*) by using experimentally evolved strains of bacteria in controlled invasions. Based on our results it was clear that invasions were promoted under fluctuating environments and if communities had not adapted to the fluctuating environment. However, the role of invader evolution on invasions was less clear, and was dependent on community evolution and/or environment during the invasion.

Throughout the experiment, environmental fluctuations played a major role in facilitating invasions. However, interestingly especially at the early stages (three and six days after invasion, Fig. 2a, b) of invasion, in the constant environment the invasion success was very persistent, and low, regardless of the invader’s or community’s evolutionary background. Thus it seems that fluctuations reveal, and constant environments buffer, evolutionary differences of strains. In the framework of current climate change this finding suggests that invasions will be increasing due to increased fluctuations. Our paper is the first to report that thermal fluctuations are able to cause similar increases in invasions as resource level fluctuations manipulated in other studies (Burke & Grime, 1996; Davis *et al.* 2000; Davies *et al.* 2005; Melbourne *et al.* 2007; Liu *et al.* 2012).

However, it was also evident that increasing fluctuations do not treat all communities equally. Especially during the first six days of the invasions, the communities that were mal-adapted to fluctuating environments experienced the strongest invasions. This is an interesting and novel finding as most of the research has emphasized the role of invader background on invasion success (Lee & Gelembiuk, 2008; Davis, 2009; Ketola *et al.* 2013) rather than the community’s properties. This finding is akin to the anthropogenically induced adaptation to invade - hypothesis (AIAI; Hufbauer *et al.* 2012), which suggests that human altered habitats, often considered disturbed, would be more vulnerable to invading species. However, instead of invaders being pre-adapted to such conditions, it seems that fluctuations or disturbances can also operate via mal-adapted communities. Thus, if climate change brings increased fluctuations to areas with a previous history of fairly constant conditions, these areas should be more vulnerable to invasions.

We did not find that evolution in fluctuating temperature pre-adapted the invader strains to be especially good at invading when environments fluctuated (Bossdorf *et al.* 2008; Hamilton *et al.* 2015). Moreover, invaders evolved in fluctuating environments were not found to be universally good invaders, yet environmental fluctuation is one of the candidates for the evolution of invasive genotypes (Lee & Gelembiuk, 2008; Davis, 2009; Ketola *et al.* 2013). The better ability of the invaders from fluctuating environment to invade was only seen if invasion occurred in constant environments (six days after invasion) and when communities had evolved in constant environments (nine and twelve days after invasion). This suggests that any evolved ability to invade in fluctuating environments was counteracted by evolved ability to resist invasions in fluctuating environments. It is interesting that the increased invasiveness was not visible after the first three days of invasion, but only became visible 6, 9 and 12 days after the invasion. The late visibility of evolutionary differences during invasions suggests that the initial differences of invading strains were small and this difference was magnified at the later stages of invasion.

The rather weak overall role of the invader itself, and strong interactions between the community’s evolutionary background, invasion environment and invader, confirms that a large part of invasions can be understood through complex eco-evolutionary dynamics (Table 1, Burns, 2006; Facon *et al.* 2006; Mächler & Altermatt, 2012). Therefore, it is important to bear in mind that the environment and genotypes always play a role and invasions success is not solely dependent on the properties of an invader (Lee & Gelembiuk, 2008; Davis, 2009; Ketola *et al.* 2013). This could mean that the quest for identifying potential invaders could be a rather weak way to mitigate invasion threat, as invasions can be dictated by environmental conditions in interplay with a community’s adaptations to environmental conditions.

To summarize, we found a large role of environmental fluctuations in aiding invasions throughout our 12 day long experimental invasion. Moreover, during the first days, environmental fluctuations also revealed a novel effect of the evolutionary background of the community on invasion success, as communities that evolved in a constant environment were especially vulnerable to invasion in the fluctuating environment. This means that climate change may cast a two-fold disadvantage for those communities that are poorly adapted to environmental fluctuations. Throughout the experiment, environment clearly remained the biggest cause of invasions. A community’s evolution in a constant environment increased the invasion risk at the early stages, and the role of the community gradually decreased during the experiment. From our experiment it is clear that thermal fluctuations threaten native populations and communities in many ways – via ecological phenomena, by facilitating invasions in general, and in evolutionary ways via fluctuation-adapted invaders and fluctuation mal-adapted communities. Our novel experiment, manipulating all key players involved in invasions in climate change altered conditions, offers not only a test the general eco-evolutionary theories, but also reveals important insights on determinants of pathogen prevalence and invasion under climate change induced conditions.

## Acknowledgements

We thank the Biological Interactions Doctoral Programme and the University of Jyväskylä Doctoral Programme in Biological and Environmental Science (KS), Academy of Finland Projects 278751 (TK), 250248 (LL), and Centre of Excellence in Biological Interactions for funding and facilities. We also thank Emily Burdfield-Steel for language editing. Authors declare no conflicts of interests.

